# Respiratory Syncytial virus NS1 protein targets the transactivator binding domain of MED25

**DOI:** 10.1101/2021.11.19.469356

**Authors:** Vincent Basse, Jiawei Dong, Andressa Peres de Oliveira, Pierre-Olivier Vidalain, Frederic Tangy, Marie Galloux, Jean-Francois Eleouet, Christina Sizun, Monika Bajorek

**Affiliations:** Université Paris-Saclay, INRAE, UVSQ, VIM, 78350 Jouy-en-Josas, France; Institut de Chimie des Substances Naturelles, CNRS UPR 2301, Université Paris-Saclay, 1 Avenue de la Terrasse, 91190 Gif-sur-Yvette, France; Unité de Génomique Virale et Vaccination, Institut Pasteur, CNRS UMR 3569, 75015 Paris, France; CIRI, Centre International de Recherche en Infectiologie, Univ Lyon, Inserm U1111, Université Claude Bernard Lyon 1, CNRS, UMR5308, ENS de Lyon, 69007, Lyon, France

## Abstract

Respiratory syncytial virus has evolved a unique strategy to evade host immune response by coding for two non-structural proteins NS1 and NS2. Recently it was shown that in infected cells, nuclear NS1 could be involved in transcription regulation of host genes linked to innate immune response, via an interaction with chromatin and the Mediator complex. Here we identified the MED25 Mediator subunit as an NS1 interactor in a yeast two-hybrid screen. We demonstrate that NS1 directly interacts with MED25 *in vitro* and *in cellula*, and that this interaction involves the C-terminal α3 helix of NS1 and the MED25 ACID domain. More specifically we showed by NMR that the NS1 α3 sequence primarily binds to the MED25 ACID H2 face, which is a transactivation domain (TAD) binding site for transcription regulators such as ATF6α, a master regulator of ER stress response activated upon viral infection. Moreover, we found out that the NS1 α3 helix could compete with ATF6α TAD binding to MED25. This finding points to a mechanism of NS1 interfering with innate immune response by impairing recruitment by cellular TADs of the Mediator via MED25 and hence transcription of specific genes by RNA polymerase II.

**Importance:** Human RSV is the leading cause of infantile bronchiolitis in the world and one of the major causes of childhood deaths in resource-poor settings. It is a major unmet target for vaccines and anti-viral drugs. RSV non-structural protein NS1 is known to antagonize the cellular immune response and was recently shown to be involved in transcription regulation of infected cells. However, the exact mechanism of this regulation is not well defined. Here we show that nuclear NS1 interacts directly with the Mediator subunit MED25 and is able to compete with a cellular transcription activator, which is activated during viral infection. We hypothesize that this interaction may underlie regulation of the expression of genes involved in the innate immune response.

## Introduction

Human RSV (hRSV) is the most frequent cause of infantile bronchiolitis and pneumonia worldwide (1). In 2005 it was estimated to have caused ~34 million acute respiratory infections in children younger than 5 years and 60,000-199,000 childhood deaths worldwide (2). Severe RSV infection is a major reason for child hospitalization. The importance of RSV-associated pulmonary disease and mortality in elderly persons has also been recognized (3). Similarly, bovine RSV (bRSV) affects cattle farms and leads to economic loss due to high morbidity and mortality among calves (4, 5). Importantly, there is still no licensed vaccine for human RSV despite over six decades of attempts (6), emphasizing the need for a better understanding of RSV pathogenesis, and more particularly the mechanisms that were developed by the virus to evade host innate immune responses.

The pathology associated with RSV infection results from both viral replication and the host immune response mediated first by the production of type I interferons (IFN-I), which induces the transcription of IFN-stimulating genes (ISG) and the production of proinflammatory mediators (7, 8). However, upon infection by RSV, IFN levels remain surprisingly low. This poor induction of IFN is attributed at least in part to the two RSV non-structural proteins, NS1 and NS2. NS1 and NS2 are unique to the *Orthopneumovirus* genus of the *Pneumoviridae* family. They diverge among the different viruses of this genus and appear to contribute to host-range restrictions (5, 9, 10). Both NS1 and NS2 also act as IFN antagonists, and many of their cytosolic targets have been identified (11, 12). As an example, NS1 inhibits RIG-I activity by interacting with MAVS as well as with TRIM25, the E3 ligase of RIG-I (13, 14). NS1 and NS2 were localized to the mitochondria (15), where they form a viral degradasome, leading to degradation of multiple target proteins, notably involved in type I IFN pathway (11). NS1 was found in the cytosol as well as in the nucleus, where it is expected to interfere with host gene expression (15, 16). In a very recent publication, NS1 was shown to associate with chromatin in promotor and enhancer regions of genes related to innate immune response to viral infection (16). By targeting these DNA regulatory regions, NS1 was suggested to suppress transcription of these genes, thus antagonizing the immune response (16). However, the exact molecular mechanism of this suppression is not well defined yet. Further study of NS1 interaction with nuclear host factors will enable a better understanding of how RSV modulates host transcription.

Based on comparison of X-ray crystallographic structures, hRSV NS1 was proposed to be a structural paralog of the hRSV matrix (M) protein (17–19). NS1 displays striking structural similarity with the N-terminal domain of the M protein, as both contain a 7-stranded β-sandwich clamped by an α-helix. In contrast to M, NS1 lacks a similar C-terminal domain but contains an additional C-terminal α-helix, α3 (Fig. 1A). NS1 α3 helix was specifically shown to be involved in the modulation of host responses (18). Mutations in the NS1 α3 helix negatively affected the transcriptional regulation of genes involved in key signalling pathways such as IFN induction and oxidative stress, resulting in 2-fold reduction of RSV replication (18).

**Figure 1:**
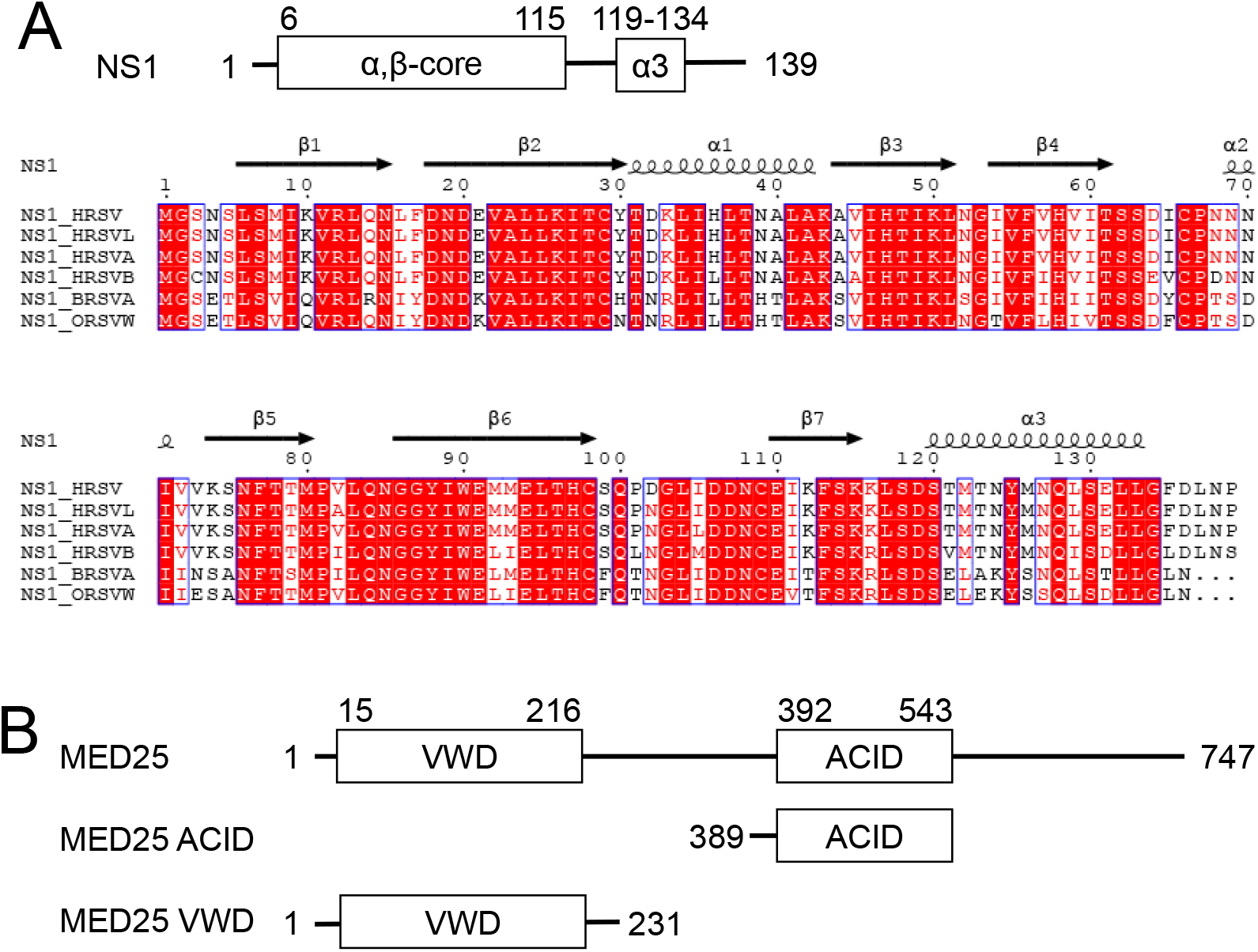
Representation of hRSV NS1 and MED25 structural organisation. **(A)** Structural organization of hRSV NS1 protein, which displays an α,β-core domain and a C-terminal α3 helix. Sequence alignment of *Orthopneumovirus* NS1 proteins: hRSV NS1 construct used in the present study, human RSV A (Uniprot P0DOE9), B (O42083) and Long (Q86306) strains, bovine RSV (Q65694) and ovine RSV (Q65703). Alignment was generated on the ClustalW server. The secondary structure elements observed in the crystallographic structure of hRSV NS1 (18) are indicated above the sequence. **(B)** Domain architecture of the Mediator subunit MED25 that contains two folded domains: the N-terminal von Willebrand domain (VWD) and the central activator interaction domain (ACID) (26, 27). The boundaries of the constructs used in this study are indicated.

Two different interactome studies of RSV NS1 pointed to an interaction of NS1 with the Mediator complex (16, 20). The Mediator complex is a nuclear multi-subunit complex that is part of the preinitiation complex required for RNA polymerase II transcription, and is a known regulator of many innate immune response genes (21–23). Several Mediator subunits were identified as potential interactors of NS1, among which MED25 (16, 20). MED25 was shown to be targeted by viral activator proteins, such as Herpes simplex virus transactivator protein VP16, which activates viral immediate-early genes during infection (24, 25). MED25 contains two folded domains: the N-terminal von Willebrand domain (residues 15-216, VWD) and the central Activator Interacting Domain (residues 392-543, ACID) (Fig. 1B). The interdomain and C-terminal regions are likely highly disordered. The ACID structure was solved by NMR and shown to be the target of both transactivation domains (TADs) of VP16 (24, 26, 27). A cryo-EM structure of the entire mammalian Mediator complex confirmed the location of MED25 in the tail module, with VWD well integrated in the tail (28) and ACID extending outside of the complex.

The putative interaction between NS1 and Mediator complex suggested by interactome studies has not been investigated so far. Having identified MED25 as an NS1 interactor in a yeast two-hybrid screen, we investigated this interaction in more details. We demonstrated that NS1 α3 helix directly interacts with the MED25 ACID domain in cells and *in vitro,* and that the residues found to be critical for innate immune response gene regulation (16) are also critical for this interaction. Moreover, we found out that NS1 α3 targets the MED25 ACID H2-face, which is the binding site of a number of TADs of transcription regulators (26, 27, 29–31). We revealed that NS1 α3 could compete with the TAD of ATF6α, involved in the innate immune response to viral infections. In contrast to transcription regulators like VP16, the small one-domain NS1 does not appear to have a distinct DNA-binding domain, and no specific DNA-binding region has been identified yet. Altogether, our results thus strongly suggest that NS1 could interfere with the host innate immune response by binding to MED25 and hindering the recruitment of transcription regulators to the Mediator, thus impairing transcription of specific host genes by RNA polymerase II.

## Results

### Identification of MED25 as a potential interaction partner of RSV NS1 by a yeast two-hybrid screen

To identify human proteins interacting with RSV NS1, we performed a yeast two-hybrid (Y2H) screen. The RSV NS1 protein fused with the GAL4 DNA binding domain (GAL4-BD) was expressed in yeast and used as a bait against prey proteins expressed from a human spleen cDNA library and fused to the GAL4 activation domain (GAL4-AD). Fifteen potential interactors of RSV NS1 were identified. We chose to focus on the MED25 subunit of the human Mediator complex, which was one of the most abundant interactors in our screen and was also identified in a previous proteomics study of host targets interacting with RSV NS1 (16, 20). In total, 10 over 156 positive yeast colonies expressed MED25. Although six of the cDNA clones expressed full-length MED25, two started at position 261 and two others at position 308. As the four cDNA clones coding for a truncated MED25 version contained the ACID domain (Fig. 1A), this strongly suggested a role of this domain in the interaction of MED25 with RSV NS1.

### NS1 interacts through its C-terminal α3 helix with MED25 ACID in cells

In order to confirm the NS1-MED25 interaction found by Y2H screening, we studied whether NS1 could interact with MED25 in cells. For that purpose, we used a split-luciferase complementation assay based on the NanoLuc enzyme (32). In this system, the 114 or the 11S NanoLuc fragments are fused to the C or N-terminus of each protein partner. To investigate the NS1-MED25 interaction, combinations of two constructs were transfected into 293T cells. Cells were lysed 24 h post transfection, luciferase substrate was added, and the luminescence, which directly depends on the interaction, was measured.

We used the RSV phosphoprotein (P), which is known to form tetramers (33–38), as a positive control. As shown in Fig. 2A, co-transfection of P-114 and P-11S resulted in a high luminescence signal, indicating a strong P/P interaction, as expected. We then used the NS1-NS1 interaction as an additional control. Although the predominant form of NS1 was reported to be monomeric (18), NS1 is also known to form dimers and higher order oligomers (15, 39). We therefore tested the NS1-114/NS1-11S pair and obtained a strong luminescence signal, revealing the capacity of NS1 to self-associate (Fig. 2A). We then tested the interaction between hRSV NS1 and full-length MED25 (Fig. 2A). When NS1-114 was co-expressed with 11S-MED25, the luminescence signal was high, confirming the interaction in cells. We then separately tested MED25 VWD and ACID domains to identify the domain involved in NS1 interaction. Transfecting NS1-114 with 11S-MED25 ACID resulted in comparable signal to that with 11S-MED25, while transfecting NS1-114 with 11S-MED25 VWD produced only background luminescence, suggesting that ACID domain was the NS1 binding domain.

**Figure 2:**
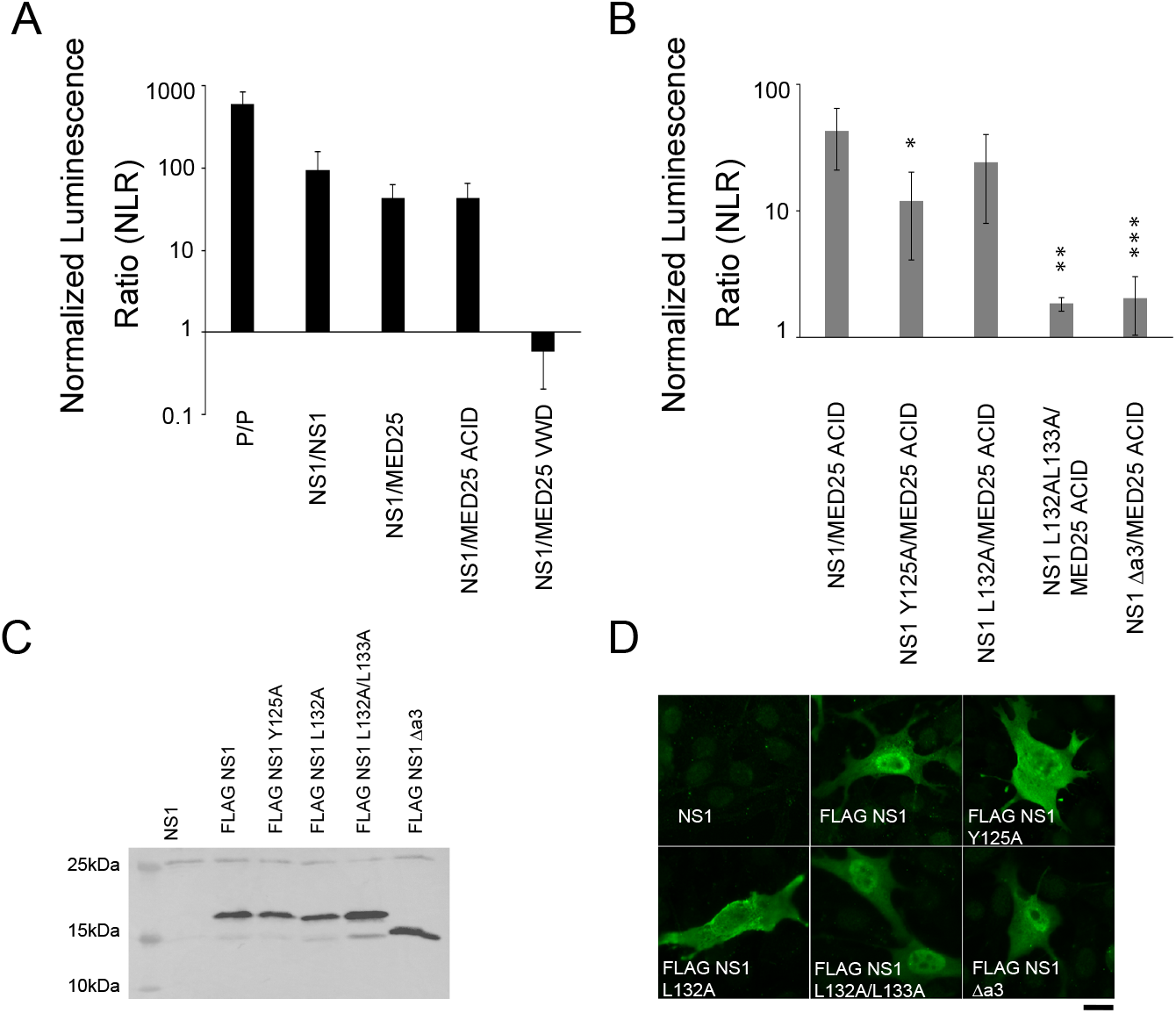
NS1 interacts with MED25 in cells. MED25 and NS1 interactions were measured using the NanoLuc assay **(A)** using MED25 domain deletions or **(B)** using MED25 ACID and FLAG NS1 WT and α3 helix mutants. 293T cells were transfected with pairs of constructs, combined as shown in the graph. P/P and NS1/NS1 were used as positive controls. The NLR is the ratio between actual read and negative controls (each protein with the empty NanoLUC vector). The graph is representative of four independent experiments, each done in three technical repeats. Data represents the means and error bars represent standard deviation across 4 independent biological replicates. *p<0.05, **p<0.01, ***p<0.001 (unpaired two-tailed t-test). **(C)** 293T cells were transfected with plasmids encoding NS1, FLAG NS1 or FLAG NS1 mutants fused to 114 NanoLUC subunit, and cell lysates were then subjected to Western analysis using anti-FLAG antibody. Size markers are shown on the left side of the gel. **(D)** BEAS-2B cells were transfected with plasmids encoding NS1, FLAG NS1 or FLAG NS1 mutants fused to 114 NanoLUC subunit. Cells were fixed, and immunostained with anti-FLAG (green) antibody followed by Alexa Fluor secondary antibody, and were analysed by microscopy. Scale bars represent 10μm.

We next asked whether the NS1 C-terminal α3 helix could be critical for the interaction with MED25 ACID. Previously, mutations in the α3 helix were shown to negatively affect transcription of key innate immune genes (18). We thus generated the same mutants: three NS1 mutants with substitutions inside the α3 helix, Y125A, L132A, and L132A/L133A, and a deletion mutant Δα3, where the α3 helix was removed. Of note, the mutants Y125A and L132A/L133A were previously shown to preserve the structural integrity of NS1 (18). Luminescence was measured in cells transfected with WT or mutant NS1-114 together with 11S-MED25 ACID (Fig. 2B). NS1 L132A/MED25 ACID co-transfection resulted in luminescence signal comparable to NS1/MED25 ACID. In contrast, co-transfection of NS1 Y125A, L132A/L133A or Δα3 with MED25 ACID significantly reduced luminescence, indicating loss of interaction. All NS1 constructs were expressed in comparable amounts in cells, as assessed by Western blot using a FLAG tag (Fig. 2C). Our results with the split-NanoLuc assay thus confirmed the NS1-MED25 interaction, and allowed to identify the MED25 ACID domain and the NS1 α3 helix as interaction domains.

Last, as MED25 has been reported to localize to the nucleus (24), and since NS1 was suggested to be actively transported to the nucleus by binding another cellular or viral protein (16), we investigated whether interaction with MED25 could influence the cellular localization of NS1. BEAS-2B cells were transfected to express FLAG-NS1 WT or mutant constructs, and the localization of NS1 protein was determined by immunofluorescence imaging after staining with anti-FLAG primary antibody (Fig. 2D). Untagged NS1 was used as negative staining control. FLAG-NS1 localized to the nucleus and to the cytoplasm, as previously reported (15, 16). None of the four tested NS1 mutants showed loss of nuclear localization, indicating that the NS1-MED25 interaction is not required for NS1 nuclear localization.

### NS1 interacts directly with MED25 ACID

Next, we investigated the interaction between human NS1 and MED25 ACID *in vitro* by GST-pulldowns using recombinant proteins. GST, GST-NS1 and GST-NS1α3 (residues 115-139) were co-expressed with MED25 ACID in *E.coli*. Bacteria lysates were incubated with glutathione beads, washed extensively and the bound complexes were analysed by SDS-PAGE and Coomassie blue staining. As shown in Fig. 3, MED25 ACID was pulled down by GST-NS1 as well as GST-NS1α3. Spurious binding was observed with GST without NS1. However, the relative band intensities between GST and retained MED25 ACID were significantly lower than with GST-NS1 or GST-NS1α3. In conclusion, our results showed that the NS1/MED25 ACID interaction is direct and mediated by the C-terminal α3 helix of NS1.

**Figure 3:**
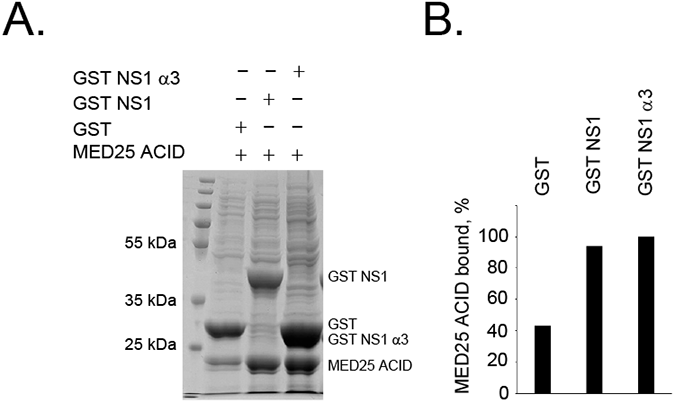
Validation of NS1-MED25 ACID interaction by GST pull-down assay. **(A)** MED25 ACID was co-expressed together with GST, GST-NS1, GST-NS1 α3 helix in *E. coli* BL21(DE3) bacteria. Bacteria lysates were clarified and the soluble proteins complexes were purified on glutathione-Sepharose beads. After extensive washing the binding of MED25 ACID to GST, GST-NS1 and GST-NS1 α3 helix was analysed by SDS-PAGE and Coomassie bleu staining. **(B)** Band intensities were quantified with J imager.

### Mapping of NS1 interaction regions on MED25 ACID by NMR

To map more precisely the NS1 α3 helix interaction site on MED25 ACID, we performed NMR interaction experiments. We titrated ^15^N-labelled MED25 ACID by an N-terminally acetylated peptide, NS1α3, corresponding to the sequence of the NS1 α3 helix. At each titration point we acquired a 2D ^1^H-^15^N HSQC spectrum (Fig. 4A). The backbone chemical shifts of MED25 ACID were assigned *de novo* by measuring 3D triple resonance experiments on ^13^C^15^N-labelled MED25 ACID. During the titration, perturbations of individual MED25 ACID amide signals were observed, showing that the NS1α3 peptide binds to MED25 ACID (Fig. 4A). For most of these residues, saturation was reached at a molar protein:peptide ratio r ~2. Due to the small size of the peptide, no significant line broadening was observed for the NMR signals of the complex as compared to free protein, which facilitated data analysis. Most perturbed signals exhibited a linear variation of chemical shifts up to saturation, indicative of a fast chemical exchange regime. Several residues, like Gly524, exhibited line broadening during titration, i.e. an intermediate exchange regime between free and bound forms (inset in Fig. 4A). The broadened signals were recovered at r ~2. Chemical shifts perturbations (CSPs, Fig. 4B) at r = 1.1 were mapped onto the 3D structure of MED25 ACID (Fig. 4C). All these perturbations were predominantly located on the H2 face of MED25 ACID, corresponding to the binding surface of the second transactivation domain (TAD2) of VP16 (26, 27, 29). Mapping of residues in intermediate exchange onto the MED25 ACID structure revealed that they also belong to the H2 face (Fig. 4D), suggesting that they report on the same binding event as those in fast exchange. An exchange rate between free and bound states of ~500 s^−1^ was estimated from the resonance frequency difference in the intermediate exchange. Intriguingly, the area perturbed by NS1α3 extends to the junction between the H1 and H2 faces, suggesting that NS1α3 binding may be accompanied by conformational rearrangement of MED25 ACID, for example by repositioning of the C-terminal α3 helix with respect to the β-barrel (Fig. 4C and 4D).

**Figure 4:**
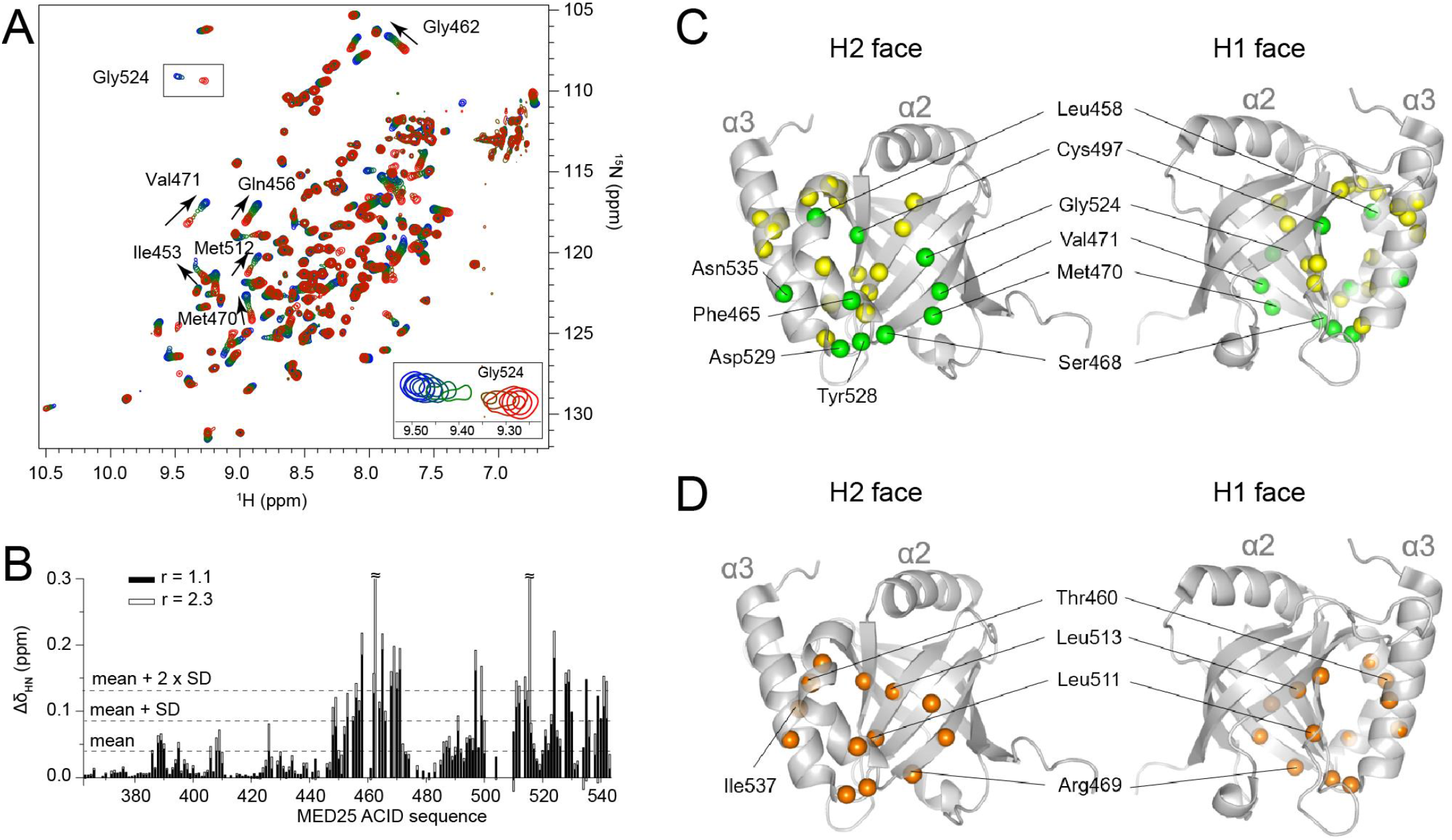
Interaction of NS1 α3 peptide with MED25 ACID followed by NMR. **(A)** Overlay of 2D ^1^H-^15^N HSQC spectra acquired during a titration of 225 μM ^15^N-labeled MED25 ACID with increasing amounts of NS1 α3 peptide. The reference spectrum without peptide is shown in red. The titration endpoint at a peptide:protein molar ratio r = 2.3 is in medium blue. Intermediate titration points at r = 0.1, 0.2, 0.4, 0.6, 0.85, 1.1, 1.4, and 1.7 are colour coded from dark orange to dark blue. Arrows show the titration direction. **(B)** Combined ^1^H and ^15^N amide chemical shift perturbations (Δδ_HN_) are stack plotted as a function of the residue number in the MED25 ACID construct for r = 1.1 (black bars) and r = 2.3 (empty bars). The bars at r = 2.3 were cut for residues Thr460 and Leu513 (Δδ_HN_ > 0.3 ppm). The mean value and mean plus one and two standard deviations (SD) for r = 1.1 are indicated by broken lines. **(C)** Chemical shift perturbations at r = 1.1 are mapped onto the structure of MED25 ACID (pdb 2xnf). Amide nitrogen atoms are drawn as spheres in green for residues with Δδ_HN_ ≥ mean+2×SD and in yellow when Δδ_HN_ ≥ mean+SD. The two views, corresponding to the H2 and H1 faces of MED25 ACID, are rotated by 180°. **(D)** Several signals are broadened at intermediate titration points due to an intermediate exchange regime, as exemplified by Gly524 in the inset shown on the ^1^H-^15^N HSQC spectra in (A). Residues in intermediate exchange regime are highlighted in orange.

Dissociation constants (K_d_) were extracted from CSPs. CSPs for residues with linear trajectories like Leu452, Met470 and Met512 were well fitted with a single binding site model (Fig. 5A-B). An average value of 17 ± 8 μM was calculated from ^1^H and ^15^N CSPs larger than mean+SD (Fig. 4B). The measured affinity is lower than those reported for individual TADs binding to MED25 ACID in the 0.5-1.5 μM range (26, 30, 31). However, the affinity of full-length NS1 may be higher than that of the NS1α3 peptide, since the α3 helix is preformed in NS1. We did not observe any significant H_Ni_-H_Ni+1_ cross-peaks typical of α-helices in 2D NOESY and ROESY spectra of free NS1α3 peptide. Only residues 123-125 and 133-138, which do not display any α-helical conformation in NS1, showed weak cross-peaks, indicating that free NS1α3 peptide remains mainly unstructured.

**Figure 5:**
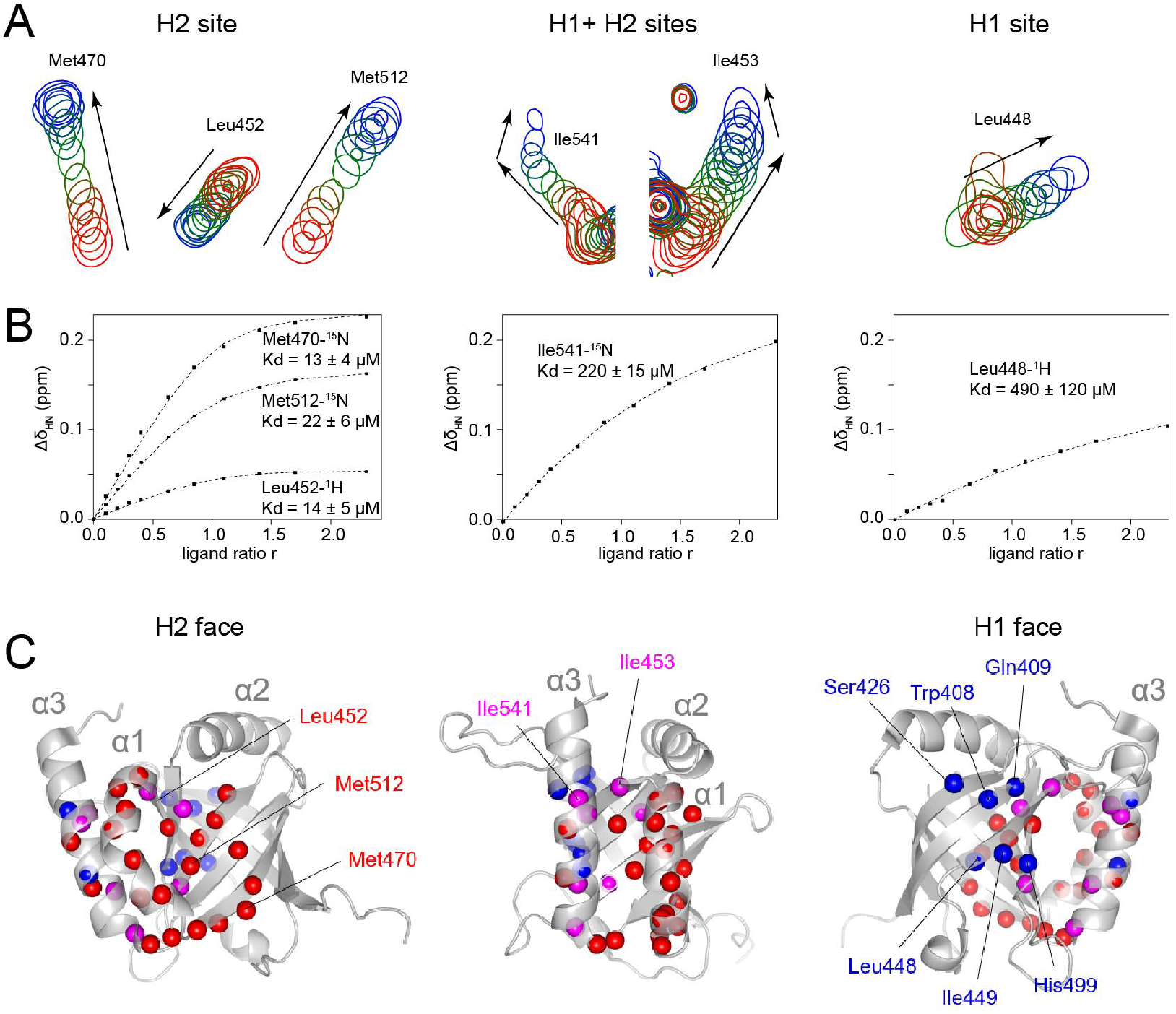
Binding modes of NS1 α3 peptide to H1 and H2 faces of MED25 ACID. **(A)** Three types of chemical shift perturbation trajectories, shown for selected residues, were observed during the ^1^H-^15^N HSQC titration of ^15^N-MED25 ACID by the NS1 α3 peptide. Arrows show the titration direction. The colour is varied from red to medium blue for molar peptide:protein ratios r = 0 to 2.3, with titration points at r = 0.1, 0.2, 0.3, 0.4, 0.6, 0.85, 1.1, 1.4, and 1.7. (B) The ^15^N or ^1^H chemical shift dimensions of the titration curves, shown in A, were fitted with a single binding site model, assuming fast chemical exchange. Experimental points are represented with solid symbols and the fitted curve in broken lines. The apparent dissociation constants K_d_ obtained from each fit are indicated. **(C)** Residues with high chemical shift perturbations are mapped onto the structure of MED25 ACID (pdb 2xnf) by representing their amide nitrogen in a sphere colour coded according to their binding mode. Residues with higher affinity, i.e. with apparent K_d_ values ranging from 7-40 μM, are indicated in red. Residues with high chemical shift perturbations, but for which saturation was not achieved at r = 2.3 are shown in blue. Residues that report on both binding events are represented in magenta. The three views are rotated by 90°.

On closer inspection, saturation was not achieved at r = 2.3 for all residues, as exemplified by Leu448 in Figure 5A. Other residues, like Ile541, displayed nonlinear chemical shift perturbation trajectories with a change at r ~1.7 (Fig. 5A). Apparent K_d_ values extracted from the binding curves were found in the 200 μM to 1 mM range (Fig. 5B). These results point to a second binding site of lower affinity. Mapping of the residues with high and low affinity onto the structure of MED25 ACID showed that residues with high affinity cluster on the H2 face, whereas residues with lower affinity cluster on the H1 face (Fig. 5C). Interestingly, the H1 face is the binding site for the TAD1 domain of VP16 (26, 27). Residues that sense the two binding modes cluster are located in between (Fig. 5C).

Taken together, our results show that the α3 region of NS1 primarily targets the H2 face of MED25 ACID, although weak binding also takes place at the H1 face. Even if the NS1α3 peptide is unstructured in its free form, unlike full-length NS1, it is expected to fold into an α-helix upon binding, like other TAD domains.

### NS1 competes with ATF6α for binding to MED25 ACID

MED25 is a target of several transcriptional activators, from cellular and viral origin (26, 27, 29, 31, 40, 41), which bind either to the H1 or H2 face of MED25 ACID through their TAD domains (30). While VP16TAD1 and the Ets family transcription factor ERM TAD were shown to bind to H1 (27, 41), VP16TAD2 and p53TAD2 bind to H2 (26, 29, 31). Previous studies have also shown that the endoplasmic reticulum stress-responsive transcription factor α (ATF6α), that functions as a master regulator of ER stress response, also targets MED25 (42, 43), and that the TAD of ATF6α (residues 40-66, Fig. 6A) binds to the H2 site (30). Since NMR results indicated that NS1 α3 helix binds to H2, we wondered whether NS1 could compete with a TAD domain, by using ATF6α. A GST-ATF6α construct containing the TAD domain (GST-ATF6αTAD, residues 1-150), bound to glutathione beads, was incubated with recombinant MED25 ACID (30 μM) and with increasing concentrations of NS1 protein (4-32 μM). The bound fractions were then analysed by SDS-PAGE and Coomassie blue staining. GST alone was used as a negative control. A truncated form of GST-ATF6αTAD was co-purified with the full form, as shown in Fig. 6B. MED25 ACID was pulled down by GST-ATF6αTAD, but not by GST (Fig. 6C), as previously published (42, 43). Adding NS1 inhibited MED25 ACID binding to GST-ATF6αTAD in a dose dependent manner (Fig. 6D, upper panel). SDS-PAGE analysis of the unbound fractions showed that increasing concentration of NS1 resulted in increasing amounts of unbound MED25 ACID (Fig. 6D lower panel). In summary, these data suggest that NS1 is able to compete for MED25 ACID binding with an H2-binding TAD domain such as ATF6αTAD.

**Figure 6:**
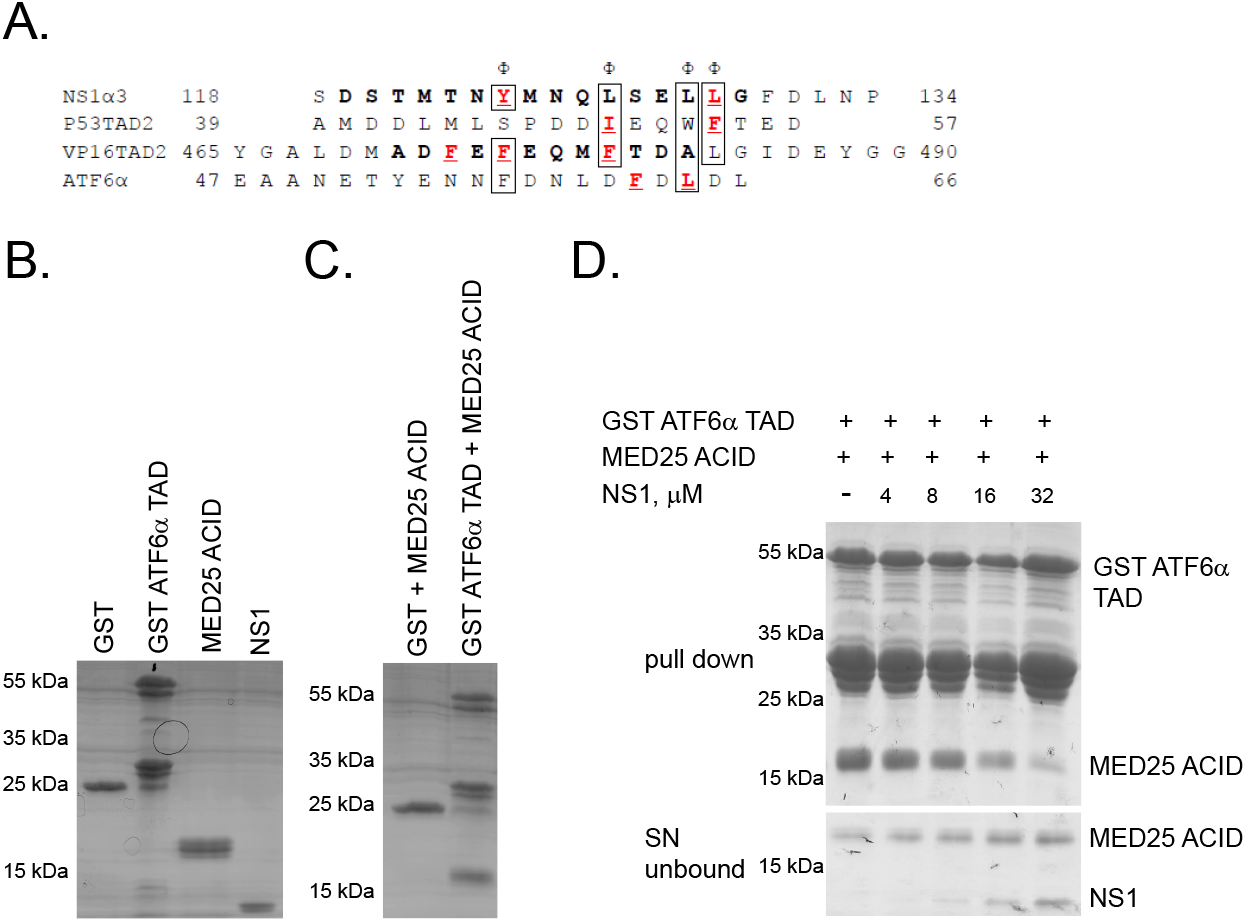
NS1 competes with ATF6α for binding to MED25. **(A)** Sequence alignment of NS1 α3 with transactivator domains of transcription factors p53, VP16 and ATF6α. The letter Φ indicates hydrophobic or aromatic amino acids. Bold letters indicate residues that form an α-helix either in the unbound state or when bound to MED25 ACID (18, 26, 31, 43). Underlined letters indicate residues that are critical for the interaction of NS1 α3, p53TAD2 and ATF6α with MED25 ACID or critical for transcription in yeast for VP16H2. **(B)** SDS-PAGE and Coomassie blue staining of purified recombinant GST, GST-ATF6αTAD, MED25 ACID and NS1 protein. **(C)** GST or GST-ATF6αTAD were expressed in *E. coli* BL21(DE3), purified on glutathione-Sepharose beads, and incubated in the presence of MED25 ACID. After extensive washing the binding of MED25 ACID to GST and GST-ATF6αTAD was determined by SDS-PAGE and Coomassie bleu staining. **(D)** GST-ATF6αTAD protein was purified on glutathione-Sepharose beads and incubated in the presence of MED25 ACID (30μM) or MED25 ACID with increasing concentration of NS1 as indicated. After incubation supernatants were collected; beads were extensively washed and the binding of MED25 ACID to ATF6αTAD and the proteins in the supernatants were analysed by SDS-PAGE and Coomassie blue staining (upper and lower gels respectively).

## Discussion

### NS1 interacts with MED25 in cells

A previous proteomics study aiming to identify host partners of RSV NS1 identified several proteins involved in transcription regulation, among them Mediator complex proteins (20). Recent NS1 co-immunoprecipitation and mass spectrometry analysis also identified subunits of the complex, among them MED25 (16). By using a Y2H screen, we identified MED25 as an interacting partner of NS1. Our NanoLuc interaction assay confirmed the NS1-MED25 interaction in cells and identified the MED25 ACID and NS1 C-terminal α3 helix as interaction domains (Fig. 2A and B). NS1 α3 was further confirmed by GST pull-down (Fig. 3) and by NMR (Fig. 4 and Fig. 5) to directly interact with MED25 ACID.

Interestingly, NS1 α3 helix was previously shown to contribute to the modulation of host response to RSV infection (16, 18). Mutation of residues Y125, and L132/L133 in the NS1 α3 helix or truncation of the entire α3 helix impacted the ability of NS1 to inhibit type I IFN. Moreover, recombinant RSV viruses carrying these mutations showed attenuated replication in IFN-competent cells and differential gene expression in the IFN pathways as compared to WT RSV (18). The same amino acids (Y125, L133) appeared to be critical for MED25 ACID interaction. Importantly these NS1 α3 helix point mutants can still properly localize to the nucleus (Fig. 2D), showing that the NS1-MED25 interaction is not required for NS1 nuclear transport. The structural integrity of these mutants has been verified previously (18). Strikingly, in the dimeric crystal structure of NS1, L133 and Y125 make intra-protomer and inter-protomer contacts, respectively, while L132 makes inter-and intra-protomer contacts, suggesting that they are buried in the structure and not available for interactions. If NS1 is monomeric, Y125 becomes accessible, whereas L132 and L133 anchor the α3 helix to the α,β-core of NS1. Since L133 appears to be critical for targeting MED25, this raises the question whether the α3 helix may dissociate from the α,β-core in solution.

Our NMR data indicate that the NS1α3 peptide preferentially binds to the H2 face of MED25 ACID, like several TADs of transcriptional regulators (26, 29–31). Our NMR titration experiment displayed similar features to those reported for TADs of transcription regulator VP16 and p53, i.e. similar concentrations to reach saturation and fast chemical exchange, suggesting similar binding modes and affinities. The 10-20 μM K_d_ obtained by NMR for NS1 α3 is indeed comparable to the 8μM value measured for the TAD2 domain of p53 by ITC (31). The 8-fold molar excess of peptide needed to reach saturation in the NMR titration by VP16-H2 H2 (26) also suggests 1-10 μM affinity. Surprisingly ATF6α (residues 40-66) binding, measured by fluorescence anisotropy, was stronger with a K_d_ of 0.5 μM (30). Comparing the sequences of the three H2-binding TADs with that of NS1 α3 did not reveal striking sequence similarity (Fig. 6A). Even residues that are critical for binding to MED25 ACID or function related to MED25 do not display any common pattern, apart from the requirement for hydrophobic residues (Fig. 6A). This is rather intriguing, but might underline that binding occurs in a multi-step process, with specificities for each TAD, as already pointed out by Henderson et al (30).

### NS1 competes with cellular TADs for targeting MED25

Transcription activator ATF6α functions as a master regulator of ER stress response. In response to ER stress, ATF6α translocates to the Golgi, where it is processed, followed by transport to the nucleus, where it activates the unfolded protein response (UPR) genes (44, 45). ATF6α was shown to recruit the Mediator complex by binding directly to the MED25 subunit (43), via the H2 site on MED25 ACID (30). Our NMR analysis showed direct binding of NS1 to the MED25 ACID H2 site (Fig. 4), suggesting that NS1 might be able to compete with ATF6α for binding to MED25. Our competition studies showed a decrease of MED25 ACID bound to ATF6α in the presence of NS1 (Fig. 6), favouring this hypothesis. Very recently it was shown that RSV infection activates the UPR, partly by activating ATF6α, to enhance virus production (46). While our study suggests that RSV could de-activate ATF6α by NS1 competing for MED25 binding, it is possible that activating and de-activating ATF6α needs to be carefully balanced during RSV infection. Even as viruses utilize the host UPR to enhance virus production and host cell survival, the invoked UPR in turn has the potential to sense viral infection and trigger anti-viral responses (47).

Very recently, NS1 was shown to associate with chromatin, and gene regulatory elements such as enhancers of genes differentially expressed during RSV infection were singled out, suggesting a new role for NS1 in regulating host gene transcription (16). Importantly, 43% of NS1 peaks identified by Chip-seq analysis coincided with Mediator peaks (16). Our results show a direct interaction between NS1 and MED25 via the H2 face of MED25 ACID, which rationalized NS1 association with Mediator peaks (16). Moreover, the Chip-seq analysis showed that the NS1 α3 helix mutant Y125A did not impact NS1 binding to chromatin, but modulated gene expression, which suggested that α3 helix may be important for interaction with a cellular partner regulating host transcription (16).

MED25 has recently emerged as one of the most significant targets for functional interactions with a range of transcriptional activators, including Herpes simplex virus transactivation protein VP16 (27, 29), ATF6α (43), ERM transcription factor (41), and p53 (31). Cellular and viral transcriptional activators that target MED25 are multi-domain proteins, which contain at least one transactivation domain (TAD) that binds the transactivator Mediator subunit MED25 and a DNA-binding domain that recognizes specific promoters/signals on target genes, which are then transcribed by RNA Pol II. Our results suggest that NS1 possesses a TAD domain and that this TAD is able to displace those of other regulation factors from the Mediator complex, thereby reducing related activation. Moreover, RSV NS1 and NS2 are the most abundantly transcribed RSV genes (15). On this basis we propose that NS1 could act as a transcription suppressor. This would present a new mechanism to control the host response upon RSV infection by interfering with activation of innate immune response genes by cellular transcriptional activators. Given the central role of NS1 in antagonizing the innate immune response to RSV, and MED25 being targetable by allosteric small molecules (30), our data could open a new avenue for RSV drug design.

## MATERIALS AND METHODS

### Plasmid constructs

Custom synthesized pciNanoLuc 114 and 11S vectors (GeneCust) were used to clone the codon-optimized hRSV NS1 and MED25 constructs using standard PCR, digestion and ligation techniques. pcineo NS1 single site mutants in the full-length construct were generated by using the Q5 site-directed mutagenesis kit (New England BioLabs), following the manufacturer recommendations. pGEX4T3 was used to clone NS1 using standard PCR, digestion and ligation techniques. pGEX NS1 α3, and pGEX NS1 single site mutants were re-cloned from pcineo vector using standard PCR, digestion and ligation techniques. MED25 (Addgene) deletion mutants were obtained by introducing start and stop codons at the appropriate site in the coding sequence (MED25 VWD aa 1-231 and MED25 ACID aa 389-543). pet41s GST ATF6*α*TAD (aa 1-150) (Addgene) contained the TAD domain.

### Y2H screen

The Y2H screen was performed as previously described (48). The DNA sequence encoding RSV NS1 was cloned by *in vitro* recombination (Gateway technology; Invitrogen) from pDONR207 into the Y2H vector pPC97-GW to be expressed as a fusion protein with the GAL4 DNA-binding domain (GAL4-BD). AH109 yeast cells (Clontech; Takara, Mountain View, CA, USA) were transformed with this construct using a standard lithium-acetate protocol. Screens were performed on a synthetic medium lacking histidine (-His) and supplemented with 3-amino-1,2,4-triazole (3-AT). A mating strategy was used to screen a commercial human spleen cDNA library (Invitrogen) established in the pPC86 vector to express cellular proteins in fusion downstream of the GAL4 transactivation domain (GAL4-AD). After 6 days of culture, colonies were picked and replica plated over three weeks to maintain selection and eliminate potential contaminants. cDNA inserts were amplified from positive yeast colonies using primers that hybridize within the backbone of the pPC86 vector. After sequencing of the PCR products, cellular interactors were identified by multi-parallel BLAST analysis.

### Bacteria expression and purification of recombinant proteins

MED25 ACID (residues Leu389-Asn543) was produced with an N-terminal 6xHis-tag followed by a T7 tag from a pET28-derived plasmid. *E. coli* BL21(DE3) bacteria transformed with the pET28 MED25 ACID plasmid were grown from fresh starter culture in Luria-Bertani (LB) broth at 37°C to an optical density of 0.6 at 600 nm, followed by induction with 0.2 mM isopropyl-β-D-thiogalactoside (IPTG) for 18 h at 20°C. Cells were lysed by sonication (4 times for 20 s each time) and lysozyme (1 mg/ml; Sigma-Aldrich) in 50 mM Na phosphate, 300 mM NaCl, 10 mM imidazole pH 8, plus protease inhibitors (Roche). Lysates were clarified by centrifugation (23,425 g, 30 min, 4°C), and the soluble MED25 ACID protein was purified on 1 ml beads loaded with Ni-NTA (GE Healthcare). The bound protein was washed extensively with loading buffer containing 25 mM imidazole and eluted with a 250 mM imidazole pH 8.

^15^N- and ^15^N^13^C-labelled MED25 ACID samples were produced in minimal M9 medium supplemented with 2 mM MgSO_4_, 100 μM CaCl_2_, 1X MEM vitamin solution (Gibco), 30 μg·mL^−1^ kanamycin, 1 g·L^−1 15^NH_4_Cl (Eurisotop, France) and 4 g·L^−1^ glucose or 3 g·L^−1 13^C-glucose (Eurisotop, France). Expression was induced with 0.1 mM IPTG. Lysis, clarification and purification, using 2 mL Ni-NTA resin (ThermoFisher, France) per liter of culture, were carried out as described for unlabelled MED25 ACID. The eluted His-tagged protein was then dialyzed into 20 mM Na phosphate pH 6.5, 100 mM NaCl buffer supplemented with 0.5 mM dithiothreitol (DTT) using a 10 kDa cut-off membrane (Spectrapor). The protein samples were further purified by gel filtration on a Superdex S75 HR 10/30 column (GE Healthcare). Samples were then concentrated to ~500 μM using 10 kDa cut-off centrifugal filter units (Amicon Ultra, Millipore) and the DTT concentration raised to 5 mM. The concentration was determined by measuring the absorption at 280 nm and applying a molar extinction coefficient of 22,460 mol^−1^· cm^−1^.

For NS1 expression, *E. coli* BL21(DE3) bacteria transformed with the pGEX-NS1 plasmid were grown from fresh starter culture in LB broth at 37°C to an optical density of 0.8 at 600 nm, followed by induction with 0.5 mM IPTG for 18 h at 20°C. Cells were lysed by sonication (4 times for 20 s each time) and lysozyme (1 mg/ml; Sigma) in 20 mM Tris-HCl, 300 mM NaCl, 5% glycerol, pH 8, plus protease inhibitors (Roche). Lysates were clarified by centrifugation (23,425 g, 30 min, 4°C), and the soluble GST-NS1 was purified on 1 ml Glutathione Sepharose beads (cytiva). The bound protein was washed with 20 mM Tris-HCl, 1M NaCl, 5% glycerol, pH 8, followed by wash with 20 mM Tris-HCl, 300 mM NaCl, 5% glycerol, 5 mM 2-mercaptoethanol, pH 8. GST-NS1 beads were then washed with 20 mM Tris-HCl, 150 mM NaCl, 2.5 mM CaCl_2_, 5 mM 2-mercaptoethanol, pH 8 and incubated with Biotinylated-thrombin protease (Novagen) over night at 4C. The supernatant NS1 fraction was collected and incubated with Streptavidin agarose (Millipore) for 1 h at 4°C in order to eliminate Thrombin. Purified NS1 was then concentrated using Vivaspin columns (Sartorius).

For GST and GST-ATF6*α*TAD expression, *E. coli* BL21(DE3) bacteria transformed with the pGEX or pet41s-ATF6*α* plasmid were grown from fresh starter culture in LB broth at 37°C to an optical density of 0.5 at 600 nm, followed by induction with 1 mM IPTG for 18 h at 20°C. Cells were lysed by sonication (4 times for 20 s each time) and lysozyme (1 mg/ml; Sigma) in 50 mM Tris-HCl, 300 mM NaCl, pH 8, plus protease inhibitors (Roche). Lysates were clarified by centrifugation (23,425 g, 30 min, 4°C), and the soluble GST-ATF6*α*TAD protein was purified on 1 ml Glutathione Sepharose beads (cytiva). The bound protein was washed extensively with 50 mM Tris-HCl and 150 mM NaCl, pH 8.

### Peptide preparation

N-acetylated NS1 α3 peptide Ac-SDSTMTNYMNQLSELLGFDLNP (RSV NS1 residues Ser118-Pro139) was synthesized by GeneCust (Luxemburg) with >95% purity, as assessed by HPLC. Aliquots of 2 mg were suspended in 1 mL MQ water and dispersed by sonication. The pH was neutralized by addition of 1 M NaOH, leading to complete dissolution. The concentration was determined by measuring the absorption at 280 nm and applying a molar extinction coefficient of 1490 mol^−1^·cm^−1^. The quality of the peptide solution was assessed by NMR. Aliquots were lyophilized for the titration experiment with ^15^N-MED25 ACID.

### Pull-down experiments

To validate NS1-MED25 ACID interaction, MED25 ACID was co-expressed together with GST, GST-NS1 or GST-NS1 *α*3 helix. *E. coli* BL21(DE3) bacteria were transformed with the pet28 MED25 ACID plasmid together with empty pGEX, pGEX NS1, or pGEX NS1 *α*3 helix. Protein induction was as for MED25 ACID alone (see above). Cells were lysed by sonication (4 times for 20 s each time) and lysozyme (1 mg/ml; Sigma) in 50 mM Na Phosphate, 300 mM NaCl, pH 8, plus protease inhibitors (Roche). Lysates were clarified by centrifugation (23,425 g, 30 min, 4°C), and the soluble proteins complexes were purified on 1 ml Glutathione Sepharose beads (cytiva). Beads were washed with 50 mM Tris-HCl and 150 mM NaCl, pH 8, and the bound proteins were analysed by SDS-PAGE and Commassie staining.

### Cell culture

293T cells were maintained in Dulbecco modified Eagle medium (eurobio) supplemented with 10% fetal calf serum (FCS; eurobio), 1% L-glutamine, and 1 % penicillin streptomycin. The transformed human bronchial epithelial cell line (BEAS-2B) (ATCC CRL-9609) was maintained in RPMI 1640 medium (eurobio) supplemented with 10% fetal calf serum (FCS; eurobio), 1% L-glutamine, and 1% penicillin-streptomycin. The cells were grown at 37°C in 5% CO_2_.

### NS1-ATF6αTAD competition assay

GST and GST-ATF6αTAD were expressed in BL21 *E.coli* and purified on Glutathione beads as described above. 50 μl GST or GST-ATF6αTAD beads were incubated with 30 μM purified MED25 ACID without or with increasing concentration of NS1 protein (4-32μM) for 2 h at 4°C. After incubation, the supernatants were collected for analysis. Beads were washed with 50 mM Tris-HCl and 150 mM NaCl, pH 8, and the samples corresponding to proteins bound to beads or recovered in the supernatant were analysed by SDS-PAGE and Commassie staining.

### NanoLuc interaction assay

Constructs expressing the NanoLuc subunits 114S and 11S were used (32). 293T cells were seeded at a concentration of 3×10^4^ cells per well in 48-well plate. After 24 h, cells were co-transfected in triplicate with 0.4 μg of total DNA (0.2 μg of each plasmid) using Lipofectamine 2000 (Invitrogen). 24 h post transfection cells were washed with PBS, and lysed for 1 h in room temperature using 50 μl NanoLuc lysis buffer (Promega). NanoLuc enzymatic activity was measured using the NanoLuc substrate (Promega). For each pair of plasmids, three normalized luminescence ratios (NLRs) were calculated as follows: the luminescence activity measured in cells transfected with the two plasmids (each viral protein fused to a different NanoLuc subunit) was divided by the sum of the luminescence activities measured in both control samples (each NanoLuc fused viral protein transfected with an plasmid expressing only the NanoLucsubunit). Data represent the mean ±SD of 4 independent experiments, each done in triplicate. Luminescence was measured using Infinite 200 Pro (Tecan, Männedorf, Switzerland).

### Immunostaining and imaging

Overnight cultures of BEAS-2B cells seeded at 4 10^5^ cells/well in 6-well plates (on a 16-mm micro-cover glass for immunostaining) were transfected with pcineo plasmids (0.4 μg) carrying the RSV codon-optimised NS1 or FLAG-NS1 WT or mutant constructs using Lipofectamine 2000 (Invitrogen) according to the manufacturer’s recommendations. At 24 h post transfection cells were fixed with 4% paraformaldehyde in PBS for 10 min, blocked with 3% BSA in 0.2% Triton X-100–PBS for 10 min, and immunostained with monoclonal anti-FLAG (1:2000; Sigma) antibodies, followed by species-specific secondary antibody conjugated to Alexa Fluor 488 (1: 1,000; Invitrogen). Images were obtained using Nikon TE200 inverted microscope equipped with a Photometrics CoolSNAP ES2 camera. Images were processed using MetaVue software (Molecular Devices).

### Nuclear Magnetic Resonance (NMR) measurements

NMR measurements were performed on a Bruker Avance III NMR spectrometer operating at a magnetic field of 18.8 T (800 MHz ^1^H frequency) and equipped with a cryogenic TCI probe. All samples were prepared in 20 mM Na phosphate pH 6.5, 100 mM NaCl, 5 mM DTT buffer and contained 7.5 % ^2^H_2_O to lock the spectrometer frequency. The temperature was set to 293 K. BEST-TROSY versions of triple resonance 3D experiments (49) were acquired on ^13^C^15^N-labeled MED25 ACID (460 μM final concentration) for backbone assignment, with a 0.2 ms recycling delay: HNCO, HNCA, HN(CO)CA, CB-optimized HNCACB and HN(CO)CACB. A standard 3D ^15^N NOESY-HSQC experiment was recorded at 700 MHz on 300 μM ^15^N-labeled MED25 ACID to confirm chemical shift assignments. ^1^H chemical shifts were referenced to DSS. NMR data were processed within TopSpin 4.0 (Bruker Biospin, Wissembourg) and analysed with CcpNmr Analysis 2.4 software (50). The titration experiment of ^15^N-MED25 ACID (245 μM) by NS1α3 peptide was carried out by recording 2D ^1^H-^15^N HSQC spectra, using a BEST-TROSY sequence. At each titration point, a lyophilized peptide aliquot was added to keep the protein concentration constant at 225 μM, starting at 0.1 and ending at 2.3 molar equivalents. Combined amide ^1^H and ^15^N chemical shift perturbations Δδ_HN_ were calculated with a scaling factor of 1/10 for ^15^N, corresponding to the ratio of gyromagnetic ratios between ^15^N and ^1^H (Eq 1):

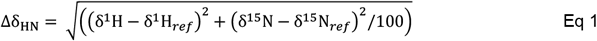

Dissociation constant Kd values were extracted by fitting MED25 ACID ^1^H and/or ^15^N chemical shift perturbations as a function of the ligand ratio, i.e. the peptide:protein molar ratio (r), with a single site binding model and assuming a fast chemical exchange regime (Eq 2), using CcpNmr Analysis software.

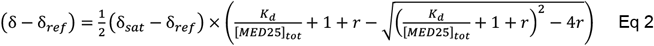

The exchange rate between free and bound states, k_ex_, was estimated from the resonance frequency difference Δν in the intermediate exchange according to k_ex_ = π * Δν.

### Illustrations

Structural representations were prepared with Pymol (Schrodinger, LLC, The PyMOL

Molecular Graphics System 1.3). Graphic rendering of sequence alignment was made with

Espript3.0 (51).

## Acknowledgements

This work was supported by Region Ile de France (DIM 1-HEALTH 2021) and Université Paris Saclay (J.D., doctoral fellowship).

## Conflicts of Interest

“The authors declare no conflict of interest.”

